# An integrative approach prioritizes the orphan GPR61 genomic region in tissue-specific regulation of chronotype

**DOI:** 10.1101/2024.11.22.624721

**Authors:** Cynthia Tchio, Jonathan Williams, Herman Taylor, Hanna Ollila, Richa Saxena

**Affiliations:** Center for Genomic Medicine, Massachusetts General Hospital and Harvard Medical School, Boston, MA; Department of Medical and Population Genetics, Broad Institute, Cambridge, MA; Department of Anesthesia, Critical Care and Pain Medicine, Massachusetts General Hospital and Harvard Medical School, Boston, MA; Department of Medicine Endocrinology, Brigham and Women’s Hospital, and Harvard Medical School, Boston, MA; Cardiovascular Research Institute; Morehouse School of Medicine; Atlanta, GA; Clinical Research Institute, Morehouse School of Medicine; Atlanta, GA; Institute for Molecular Medicine, HiLIFE, University of Helsinki, Helsinki, Finland

**Keywords:** Morning chronotype, ease of getting-up, circadian rhythm, rs12044778, *GPR61*, *AMIGO1*, *ATXN7L2*, genetic pleiotropy, knockout models

## Abstract

**Objectives:** Chronotype, a manifestation of circadian rhythms, affects morning or evening preferences and ease of getting-up. This study explores the genetic basis of morning chronotype and ease of getting-up, focusing on the G protein-coupled receptor locus, GPR61.

**Methods:** We analyzed the genetic correlation between chronotype and ease of getting-up using linkage disequilibrium score regression with summary statistics from the UK Biobank (n=453,379). We prioritized shared signals between chronotype and ease of getting-up using the Human Genetic Evidence (HuGE) score. We assessed the significance of GPR61 and the lead variant rs12044778 through colocalization and *in-silico* analyses from ENCODE, Genotype-Tissue Expression, Hi-C, and Knockout Mouse Project databases to explore potential regulatory roles of causal genes.

**Results:** We identified a strong genetic correlation (Rg=0.80, P=4.9 x10^324^) between chronotype and ease of getting-up. Twenty-three genes, including three circadian core clock components, had high HuGE scores for both traits. Lead variant rs12044778 in *GPR61* was prioritized for its high HuGE score (45) and causal pleiotropy (posterior probability=0.98). This morningness variant influenced gene expression in key tissues: decreasing *GPR61* in tibial nerve, increasing *AMIGO1* in subcutaneous adipose, and increasing *ATXN7L2* in the cerebellum. Functional knockout models showed *GPR61* knockout increased fat mass and activity, *AMIGO1* knockout increased activity, and *ATXN7L2* knockout reduced body weight without affecting activity.

**Conclusions:** Our findings reveal pleiotropic genetic factors influencing chronotype and ease of getting-up, emphasizing *GPR61*’s rs12044778 and nearby genes like *AMIGO1* and *ATXN7L2*. These insights advance understanding of circadian preferences and suggest potential therapeutic interventions.

**SIGNIFICANCE:** This study investigates the genetic underpinnings of chronotype preferences and ease of getting up, with a focus on the orphan G protein-coupled receptor GPR61 and the locus lead variant rs12044778. By combining genomic data with *in silico* functional analysis, we provide mechanistic insight into a locus for morning chronotype and ease of getting in the morning. We identified the variant rs12044778 as a key regulator of *GPR61* and nearby genes *AMIGO1* and *ATXN7L2* influencing circadian and metabolic traits. Our findings shed light on the intricate genetic networks governing circadian rhythms, suggesting potential therapeutic targets for disorders of the circadian rhythm.

## INTRODUCTION

Chronotype, a manifestation of the internal circadian clock, significantly influences individual preferences for morningness-eveningness, categorizing us as ‘morning larks’ or ‘night owls’ ^1^. This behavioral expression is not only shaped by age, gender, social obligations, and environmental inputs but is also intricately wired into the molecular mechanisms of the circadian rhythm ^2^. These mechanisms, which involve complex interactions between genetic factors and environmental cues, play a crucial role in aligning our physiological processes with the 24-hour day-night cycle ^3^.

Compelling evidence suggests that disruptions to circadian timing are closely linked to the development of diseases, with a particular emphasis on metabolic disorders ^3^. The association of chronotype with various aspects of health, such as sleep disorders, cognitive and physical performance, chronic metabolic and neurologic diseases, cancer, and premature aging, becomes particularly pronounced when there is a misalignment between one’s internal chronotype and the external environment ^4^. This mismatch increases the risk of developing diseases.

Chronotype, the individual variation in circadian rhythm preferences, plays a crucial role in determining the ease of getting up. Morning larks, characterized by an early chronotype, typically find it easier to wake up early and feel more alert during the morning hours. In contrast, night owls, with a later chronotype, may experience challenges waking up early, as their internal biological clock aligns more favorably with later parts of the day. The connection between our internal timing system and our daily wakefulness patterns is highlighted by the link between chronotype and ease of getting up; additionally, studies have indicated a genetic link to chronotype, further influencing the ease of getting up ^1,5^.

Genetic factors contribute to an individual’s predisposition towards being a morning lark or a night owl, impacting their circadian rhythm and, consequently, their waking habits. This genetic connection emphasizes the intricate interplay between biological factors and the ease of getting up, highlighting the personalized nature of our sleep-wake preferences ^5^. However, despite the recognized genetic influence on chronotype and waking habits, there is a notable gap in studies that delve into the functional aspects of specific genetic loci implicated in these traits. For instance, the chronotype study by Jones et al. ^1^ identifies 351 loci associated with chronotype, yet functional studies elucidating their roles remain scarce. Similarly, GWAS on the ease of getting up, such as those made publicly available by the Neale Lab, reveals 55 significant loci, yet analyses of these findings through resources like the Open Targets platform ^6,7^ underscore a lack of functional insights. In our current study, we aim to address this gap by using an *in silico* functional approach relevant to circadian rhythms to investigate the functionality of one of the regulatory variants, rs12044778, in *GPR61,* a G-protein coupled receptor (GPCR).

GPCRs are the largest class of membrane receptors ^8^; however, 150 out of 900 GPCRs are classified as orphans due to the lack of knowledge about their endogenous ligands, hence limiting our understanding of their biological significance ^9^. Interestingly, GPCRs are the target of 36% of FDA-approved drugs, reflecting opportunities for further research^10^. All this further increases our interest in starting with rs12044778 for functional follow-up of chronotype loci. Our overall aim is to provide a deeper understanding of the genetic underpinnings of chronotype and the ease of getting up, shedding light on the functional roles of specific genetic elements in shaping individual sleep-wake preferences. Here, we present novel findings that advance our understanding of the genetic basis of morningness chronotype and ease of getting up in the morning. Through a focused investigation of the *GPR61* gene, specifically its regulatory variant rs12044778, we have identified a significant association with other circadian rhythm traits. The combination of genome-wide association studies (GWAS) with functional analyses provides a comprehensive view of how this variant might influence chronotype and ease of getting up. Our results also highlight the intricate genetic factors contributing to chronotype and ease of getting up, offering new insights and potential directions for future research in the field of chronobiology.

## METHODS

### Study populations

We used publicly available GWAS summary statistics from the UK Biobank (UKB) Sleep-reported Traits GWAS from the Sleep Disorder Knowledge Portal ^11^. This included the UK Biobank component of the morningness chronotype GWAS (N= 453,379) from Jones and Lane et al., 2019 ^1^. Participants were asked to identify their chronotype by responding to the question: “Do you consider yourself to be?” They could choose from one of six options: “Definitely a ‘morning’ person”, “More a ‘morning’ than ‘evening’ person”, “More an ‘evening’ than a ‘morning’ person”, “Definitely an ‘evening’ person”, “Do not know”, or “Prefer not to answer”. Participants’ ease of getting up in the morning was assessed through the question: “On an average day, how easy do you find getting up in the morning?” The response options included: “Not at all easy”, “Not very easy”, “Fairly easy”, “Very easy”, “Do not know”, or “Prefer not to answer”.

### Genetic Correlation and Human Genetic Evidence

We used tools implemented in the Sleep Disorder Knowledge Portal for genetic correlation analyses and to prioritize association signals for *in-silico* functional exploration. First, we used LD score regression to calculate the genetic correlation between chronotype and ease of getting up ^12,13^. This method quantifies the extent of shared genetic architecture between the two traits. Next, to evaluate the aggregated genetic support for the involvement of genes near the signal locus, we examined the Human Genetic Evidence (HuGE) score, which quantifies genetic support for the involvement of the gene of interest in the phenotype of interest using a Bayesian approach ^14^. The common variation score reflects associations with genome-wide significance (p < 5e-8), prioritizing genes with coding variants and genes nearest to the strongest association signal.

### Colocalization Analysis

Colocalization analysis was performed using HyprColoc (Hypothesis Prioritization in Multi-Trait Colocalization) ^15^, and the default parameters prior.1=1e-4 and prior.2=1e-4 were used. This analysis focused on the 1Mb genomic region encompassing rs12044778 (Chr1:109,832,494-110,341,028; Genome Build: hg19/GRCh37). Locuszoom ^16^ was used to create regional association plots for the lead variant rs12044778 in the GPR61 gene for chronotype and ease of getting up. Using datasets from the common metabolic disease knowledge portal, we conducted a PheWAS to identify associations of rs12044778 with other circadian timing phenotypes in the sleep knowledge portal ^11^, and we used a p-value cutoff of 0.005 (0.05/10 circadian timing phenotypes).

### *In-silico* Functional Annotation

Publicly available data from Hi-C sequencing was employed to explore the 3D chromatin interactions surrounding rs12044778 in the cerebellum and adipose progenitor mesenchymal stem cells in the 3DIV database^17^. Chromatin was fixed, processed, and sequenced to produce a contact matrix, which was normalized to identify topologically associating domains (TADs) visualized as blue triangles on the heatmap. Interaction frequencies were detailed in an annotated chromosome ideogram, revealing the complex interaction landscape at the rs12044778 locus and its potential regulatory impact on nearby genes. We used HaploReg 4.2 ^18^ to explore the predicted regulatory potential of rs12044778, particularly its effect on transcription factor binding. The analysis included assessing Delta (Δ), which represents the difference in predicted binding scores or binding affinity between the reference allele and the alternate allele for a specific TF binding motif; this numerical value quantifies the change in binding strength due to genetic variation. The binding of transcription factors (EGR1, ELF1, GABPA, and NR2F2) to GPR61 at locus rs12044778 was confirmed using K562 cell ChiP-seq experiments from ENCODE (ENCFF568SWK_K562_EGR1, ENCFF246JLN_K562_ELF1, ENCFF108UGI_K562_GABPA, and ENCFF092LQW_K562_NR2F2). We used the Integrated Genome Browser (IGB) to visualize the ChiPseq peaks.

Tissue-specific eQTL analysis was performed to assess the impact of rs12044778 on the gene expression of nearby genes in various tissues using the GTEx resource v8 (n=838 total participants with data from 54 tissues). To distinguish between nearby unrelated eQTL signals and those driven by rs12044778, we assessed the co-localization of genetic association signals for morning chronotype/ ease of getting up and the eQTL signals using HyprColoc ^15^ (parameters prior.1=1e-4 and prior.2=1e-4) and generated regional association plots for the locus rs12044778 (*AMIGO1*, *ATXN7L2*, *GPR61*, and *GNAI3*) with locuszoom ^16^.

RNA-seq data from the Human Protein Atlas (HPA) was queried to analyze the gene expression levels of *GPR61*, *GNAI3*, *AMIGO1*, and *ATXN7L2* across various human tissues ^19,20^.

### Body Composition and activity in knockout mouse models from the public IMPC resource

To better understand the mechanism behind the metabolic functions of rs12044778 nearby genes, we obtained mouse knockout data from the IMPC; however, a list of 363 phenotypes was publicly available only for *GPR61*, *AMIGO1*, and *ATXN7L2* global knockout (KO) mice models ^21,22^. The IMPC is an international effort to identify the function of every protein-coding gene in the mouse genome to provide transformative insights into the genetics of diseases. The KO models were generated using the CRISPR-CAS9 technology in the C57BL/6N mice as a background, and C57BL/6N mice served as the control or wild-type (WT) model. The research resource identification initiative (RII) number for *GPR61* KO is MMRRC_047960-UCD, *AMIGO1* is MMRRC:046579-UCD, and *ATXN7L2* is MMRRC:043412-UCD. The mice were maintained in a 12-hour light-dark cycle, and testing was conducted during the light phase.

The open field test assessed the activity and exploratory behaviors of mice using Accuscan Versamax (VMX 1.4b) equipment. The mice, 9 weeks old, were left undisturbed for 30 minutes before being placed in the center of the apparatus to roam freely for 20 minutes. Results were analyzed using two-way ANOVA and Tukey post hoc test in GraphPad Prism ^21,22^.

The mice’s body composition was measured using DEXA. Each mouse, 14 weeks old, was anesthetized for the analysis. The DEXA scan focused on the whole body, excluding the head, and assessed parameters such as body weight, length, fat and bone mass, bone mass density, and lean mass. The results, expressed as mean ± SEM, were analyzed using two-way ANOVA followed by Tukey post hoc test in GraphPad Prism ^21,22^.

## RESULTS

### Genetic evidence implicates 23 genes, including *GPR61*, shared between morningness chronotype and ease of getting up in the morning

First, genetic correlation analyses using LDSC ^12,13^ between morningness chronotype and ease of getting up in the morning found a large extent of genetic overlap between the two traits (r_g_=0.80 ± 0.01, Pvalue=4.94 X 10^−324^). To identify pleiotropic loci, we next calculated the Human Genetic Evidence (HuGE) score^12^, a score that systematically aggregates both common and rare genetic variants to evaluate their collective impact on gene function and disease association. In addition to pleiotropy and shared genetic architecture, this analysis allows us to prioritize genes for functional studies ^14^. The HuGE score analysis prioritized 23 genes implicated in both traits with ‘very strong’ genetic evidence (**Figure 1**). Notably, the canonical core clock genes *PER3*, *PER2*, and *CRY1,* as well as neuropeptide receptors *HCRTR2* and *NMUR2* involved in sleep/wake regulation, were also prioritized by this approach. These genes provide proof of the principle that canonical circadian pacemaker genes contribute to both ease of getting up in the morning and to chronotype. Here, we chose to examine the potentially novel and less well-studied pleiotropic locus encompassing the orphan receptor *GPR61* with strong genetic evidence of association in morningness chronotype (HuGE =45) and ease of getting up (HuGE =45).

**Figure 1:**
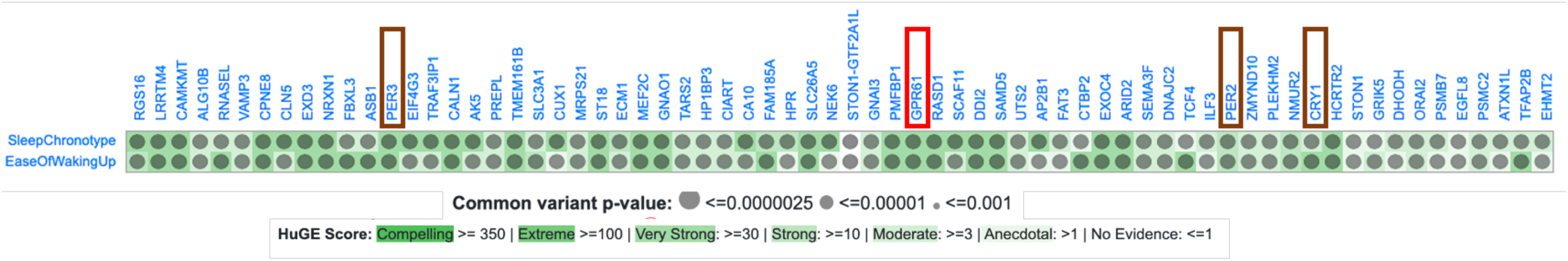
GPR61 is associated with chronotype and ease of waking up. The plot displays the Human Genetic Evidence (HuGE) score calculated to quantify the genetic support of the involvement of GPR61 (red box) in GWAS traits (SleepChronotype: Morningness Chronotype and EaseOfWakingUP: Ease of Waking up in the morning). The canonical clock genes PER3, PER2, and CRY1 are highlighted by a brown box. The score was calculated from GWAS significant loci with a p-value cutoff of 5e-8.

Regional association plots (**Figure 2a, 2b**) highlighted rs12044778 within the intron of *GPR61* as the lead variant for both chronotype and ease of getting up in the *GPR61* genomic region (Chr1:109,832,494-110,341,028; Build: GRCh37). Notably, the HuGE score of other nearby genes (*MYBPHL, SORT1, SYPL2, ATXN7L2, AMIGO1, CYB561D1, GNAI3, GNAT2, GSTM2, GSTM1, GSTM3, GSTM5, GSTM4,* and *EPS8L3*) showed no (HuGE=1) to moderate (HUGE=3) evidence, with the exception of PSMA5 (HuGE score =45 for chronotype and HuGE = 1 for ease of getting up (**Figure 2c**).

**Figure 2:**
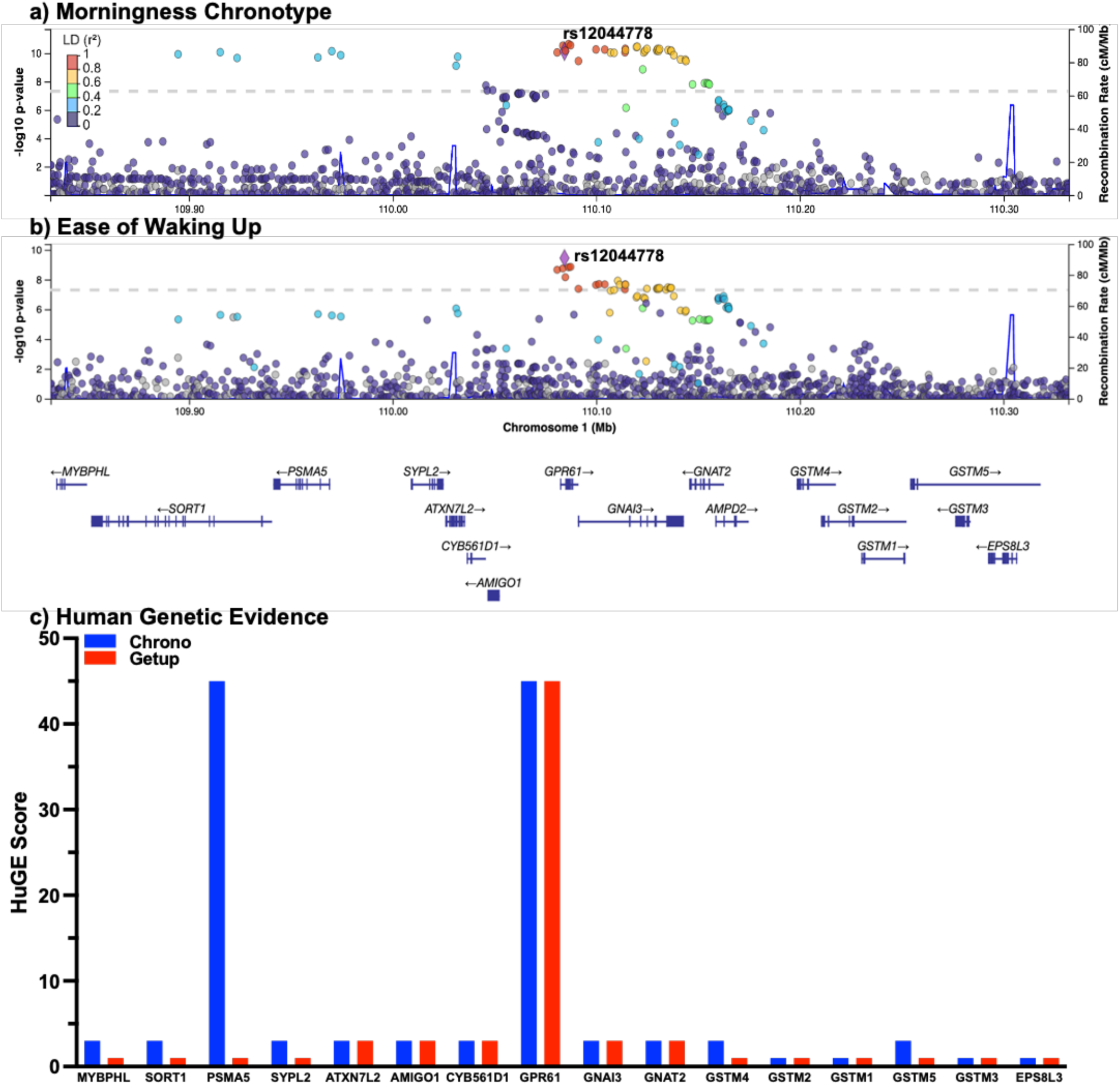
Colocalization analysis reveals a shared variant at the GPR61 locus in GWAS of morningness chronotype and ease of waking up. Regional association plot at the colocalized GPR61 locus with GWAS significance (P-value < 5e-8) for (**a**) morningness chronotype and (**b**) ease of waking up. The y-axis shows the −log10 P value for each variant in the region, and the x-axis shows the genomic position. Each variant is represented by a filled circle, with the rs12044778 lead variant colored purple and nearby variants colored according to the degree of linkage disequilibrium (r2) with rs12044778 in the 1000G EUR population. The lower panel shows genes located in the displayed region, and the blue line corresponds to the recombination rate. The Human Genetic Evidence (HuGE) score (**c**) for the genes in the genomic region of the lead variant rs12044778 Chr1:109,832,494-110,341,028 (Genome Build: hg19/GRCh37)

Colocalization analyses for both traits using HyperColoc ^15^ identified lead variant rs12044778 as the most likely causal pleiotropic variant in that region with a posterior probability (pp) of 0.9757. The rs12044778 allele A (frequency: 0.65-0.89 across Gnomad populations ^23^) was associated with morningness chronotype (β= −0.0229±0.0035 for morning chronotype; Pvalue=7.5X10^−11^) and ease of getting up (β= −0.0129±0.0020 for ease of getting up; Pvalue=3.6X10^−10^ (**Table S1**).

### *GPR61* intronic variant rs12044778 is associated with other circadian timing traits

To assess whether rs12044778 is associated with other objective or self-reported circadian phenotypes, we generated a PheWAS plot (**Figure 3**) using datasets from the sleep knowledge portal ^11^. We found that rs12044778 allele A is significantly associated with earlier most active 10-hour timing (β=0.0204±0.0062; Pvalue=0.0011) and earlier least-active 5-hour timing (β=0.0153±0.0063; Pvalue=0.014) derived from 7-day actigraphy measures in the UK Biobank that provide objective measures of sleep timing, complementing the self-reported phenotypes; the p-value threshold was set as 0.005 (0.05/10 circadian phenotypes) ^11,24^ (**Figure 3 & S1)**.

**Figure 3:**
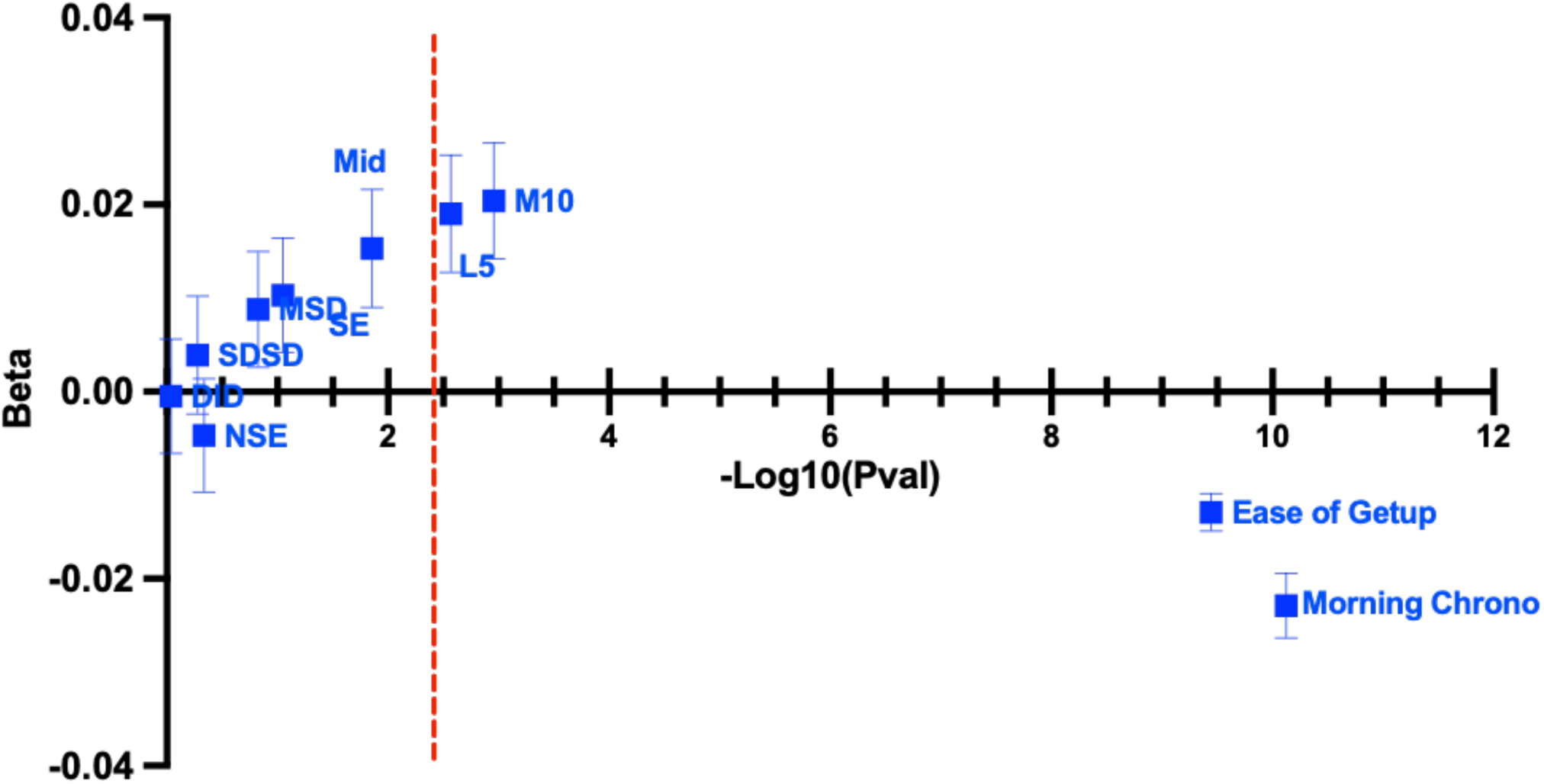
Association of the colocalized lead variant rs12044778 with accelerometer-derived sleep and activity timing measures. The plot displays the variant-level phenotypic associations for the colocalized lead variant rs12044778 expressed as effect size (Beta; y-axis) and significance (–log10(p-value)). The dashed line is the significance threshold of 0.005 (0.05/10 Circadian Phenotypes). Morning Chrono: Morningness Chronotype; Ease of Getup: Ease of waking up in the morning; M10: Most-active 10-hour timing; L5: Least-active 5-hour timing, and M5: Sleep Midpoint timing, SE: Sleep efficiency, MSD: Mean sleep duration, SDSD: Standard deviation o of sleep duration, NSE: Number of sleep episodes, DID: Diurnal inactivity duration (Table S1).

### The lead variant rs12044778 is a regulatory variant that influences the tissue-specific expression of *GPR61*

To assess whether rs12044778 affects the gene expression of nearby genes, we performed a tissue-specific expression quantitative traits loci (eQTL) analysis in subcutaneous adipose tissues and the brain tissues involved in sleep and circadian processes (**Table S3**) ^26^. In **Figure 4C**, we found that the rs12044778 morningness allele A significantly decreases the gene expression of *GPR61* in tibial nerve (Normalized Expression size (NES) = −0.35, Pvalue= 1.65e-12), increases *AMIGO1* expression in subcutaneous adipose tissue (NES=0.25, Pvalue=5.74e-7), and increases *ATXN7L2* gene expression in the cerebellar hemispheres (NES=0.34, Pvalue=1.61e-5). As expected, variants in linkage disequilibrium (r^2^ ≥ 0.80, EUR) with rs12044778 shared the same eQTL tissue-specific expression pattern (**Table S3**).

**Figure 4:**
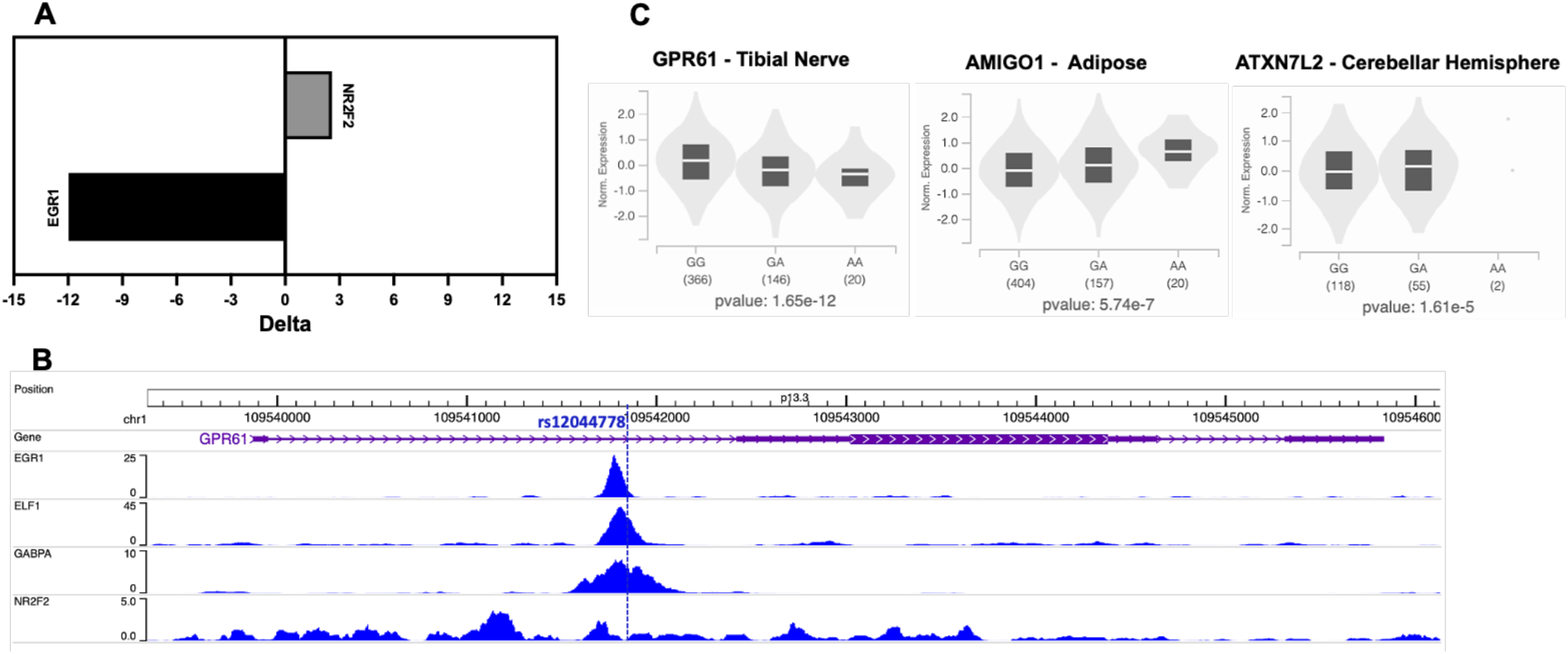
The lead variant rs12044778 in GPR61 regulates the gene expression of nearby genes. Plot (**A)** displays the transcription factors whose binding is predicted to be affected by the variant rs12044778, and it is expressed in delta, which is the binding affinity between the reference allele and the alternate allele for a specific TF binding motif. (**B**) shows rs12044778 is at the binding site of EGR-1, ELF1, GABPA, and NR2F2 transcription factors using K562 cell Chip-Seq experiment from ENCODE. Plot (**C**) is displayed as normalized gene expression and rs12044778 genotype in Gtex v.8 data with (the number of participants) for each genotype below it. The variant rs12044778 decreases the gene expression of GPR61 in nerve tibial, increases AMIGO1 expression in subcutaneous adipose tissue, and increases ATXN7L2 gene expression in the cerebellum

To differentiate between unrelated nearby eQTL signals and eQTL signals driven by rs12044778, we evaluated the colocalization of genetic association signals and eQTL signals identified above (**Table S3**). Interestingly, the lead signal rs12044778 colocalized with *ATNX7L2* expression in the cerebellar hemisphere (**Figure S2D**) for ease of getting up (pp=0.990). In the tibial nerve (**Figure S2E**), the lead signal rs12044778 colocalized with *GPR61* expression in the morningness chronotype (pp=0.9992) and ease of getting up (pp=0.9999) (**Table S4**). Moreover, the lead signal rs12044778 colocalized with *ATNX7L2* expression in the cerebellar hemisphere (**Figure S2D**) for ease of getting up (pp=0.990), suggesting a possibly broad regulatory effect from rs12044778 on local gene expression.

A possible mechanism for activating gene expression is by affecting transcription factor binding or by physical contact with the three-dimensional structure at the promoter region. To determine whether rs12044778 regulates nearby genes via chromatin loops, using 3DIV ^17^, we generate a Hi-C chromatin interaction loop plot (**Figure 5**) from the cerebellum (Experiment ID: ENCLB672PAB and ENCLB174TEA) and adipose-progenitor mesenchymal stem cells (Experiments ID: GSM1267200 and GSM1267201) ^17^. Adipose-progenitor mesenchymal stem cells are a key cell type involved in adipogenesis and broader metabolic processes. Since chronotype and circadian rhythm genes have been linked to metabolic traits ^1,24^, and GPR61 deficiency in mice leads to obesity ^25^, understanding its genetic regulation in tissues directly involved in these metabolic processes can be particularly insightful. The lead variant rs12044778 (chr1:109541786; Build: GRCh38) in *GPR61* interacts with promoters of nearby genes (*CSF1, EPS8L3, SARS1, SORT1, PSRC1, CELSR2, AHCYL1, AMPD2, GNAT2, KIAA1324, AMIGO1, GNAI3, ATXN7L2, GSTM2, GSTM1, GSTM5*) in mesenchymal stem cells (**Table S2**). Moreover, we found that rs12044778 is a regulatory variant because it was predicted to alter the binding of transcription factors EGR1 (Δ = −11.97), HOXB3 (Δ = −0.34), and NR2F2 (Δ = 2.58) (**Figure 4A**) ^18^. Using ENCODE experimental ChiP-seq data from ENCODE, we validated that rs12044778 is at the binding site of EGR-1 and NR2F2 and also for ELF-1 and GABPA transcription factors (**Figure 4B**).

**Figure 5:**
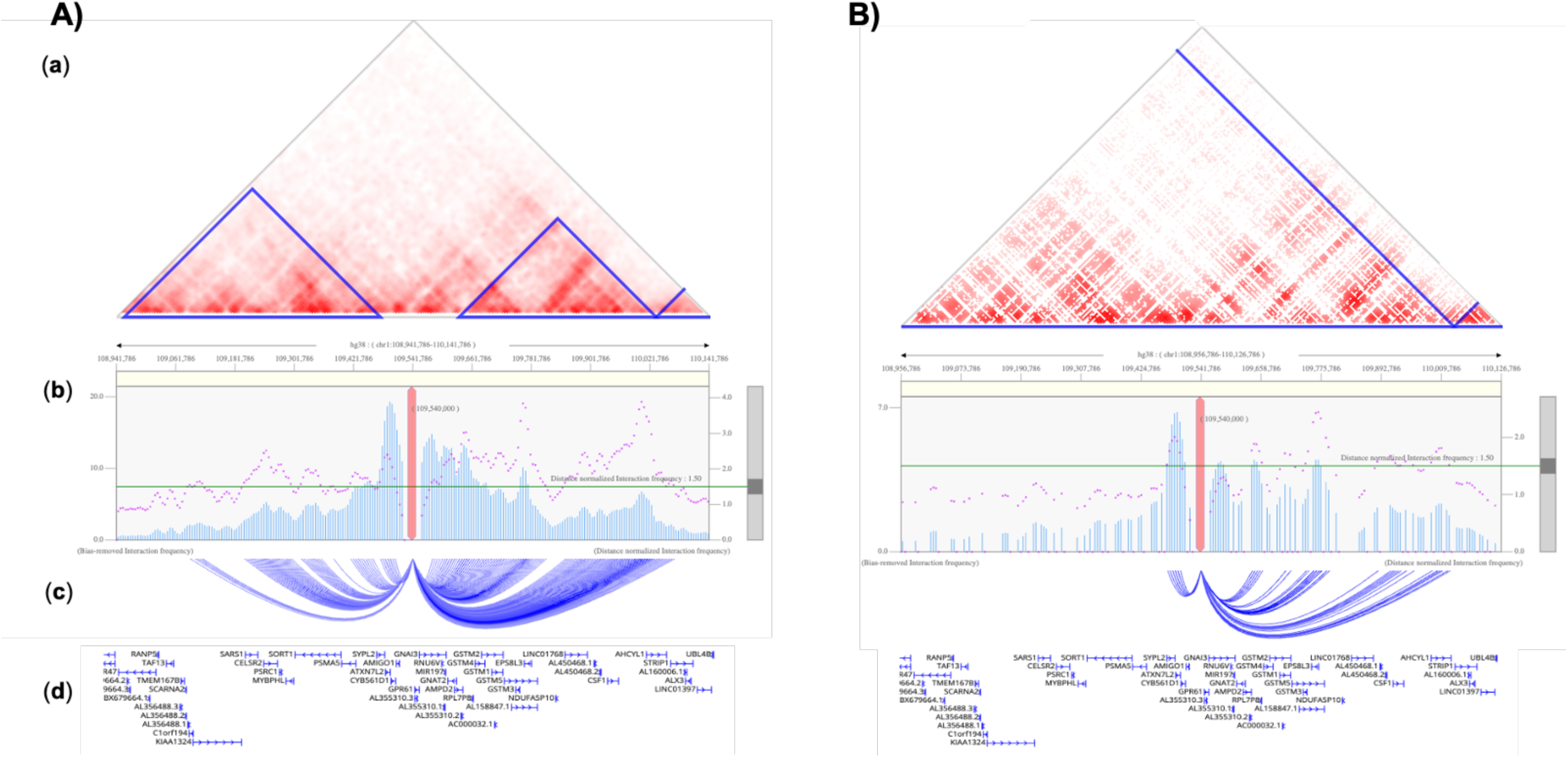
The tissue-specific interaction of the lead variant rs12044778 with nearby genes. Hi-C Interaction analysis of the rs12044778 locus in Adipose Progenitor Mesenchymal Stem Cells (A) and astrocyte of the cerebellum (B). (a) Heatmap depicting the Hi-C contact map of chromosome 1, highlighting the interaction frequencies within a specific genomic region. Topologically associating domains (TADs) are indicated by blue triangles, demarcating the structural units of chromosomal folding. (b) Bar and dot plots represent the bias-removed interaction frequencies (bars) and their corresponding distance-normalized interaction frequencies (dots), providing a detailed view of the chromatin interactions within and across the TAD boundaries.(c) Chromosome ideogram with gene annotations (d) and blue interaction arcs. The arcs visualize physical contacts between regions, elucidating the three-dimensional organization of the genome in relation to gene regulatory elements and the SNP rs12044778.

### The nearby genes regulated by rs12044778 have a unique expression pattern in the human hypothalamic regions

To investigate how nearby genes *GPR61, GNAI3, AMIGO1,* and *ATXN7L2* to rs12044778 might influence metabolic and circadian functions, we analyzed their gene expression patterns in human subcutaneous adipose, pituitary gland, cerebellum, and hypothalamus (**Figure 6A**). Dissection of the expression profile in the hypothalamic regions showed that *AMIGO1* is more enriched in the arcuate nucleus, dorsomedial hypothalamic nucleus, lateral hypothalamic area, ventromedial hypothalamic nucleus, and preoptic area (**Figure 6B**) when compared to *GPR61*, *GNAI3*, and *ATXN7L2*.

**Figure 6.**
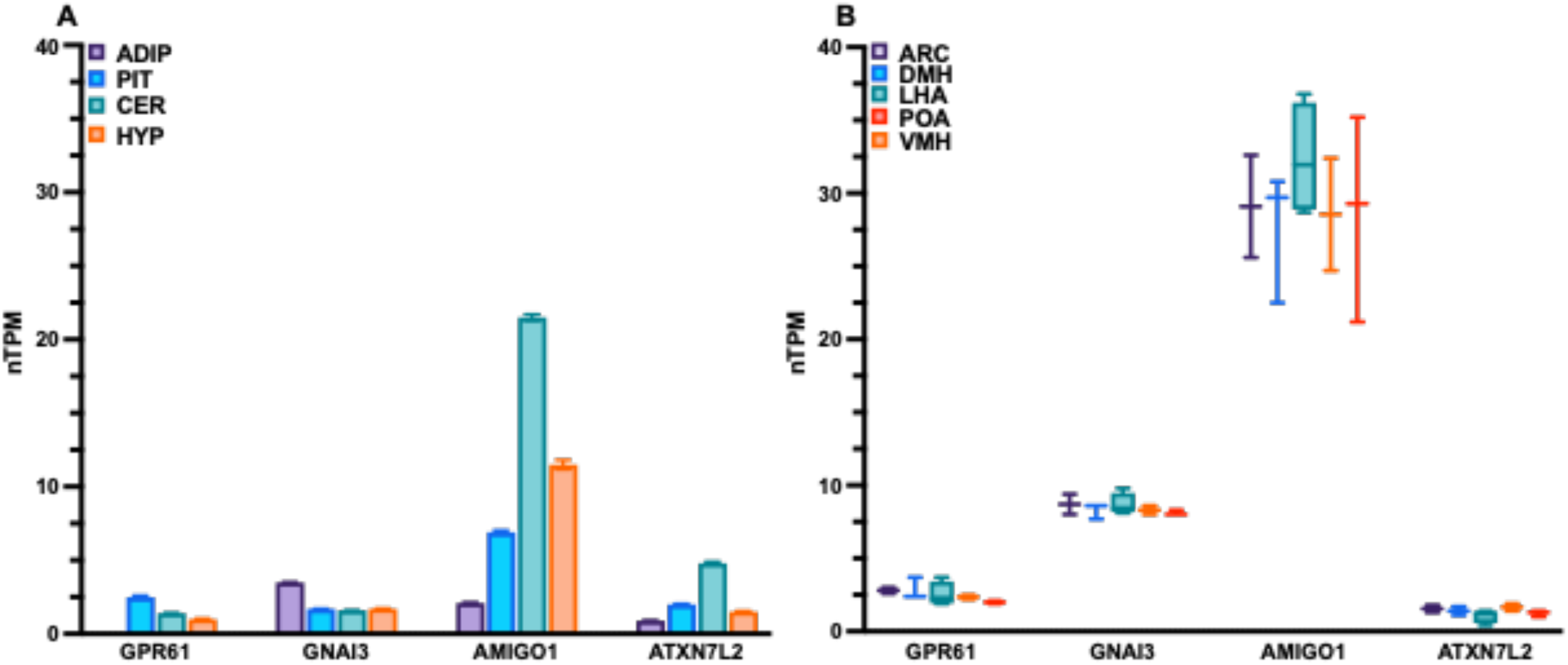
The expression profile of nearby genes regulated by rs12044778 in humans. (**A**) The bar graph is the gene expression in nTPM of *GPR61, GNAI3, AMIGO1,* and *ATXN7L2* gene expression in the subcutaneous adipose tissues (ADIP) hippocampus (HIP), pituitary gland (PIT), cerebellum (CER), and hypothalamus (HYP). (**B**) The box-whisker plot is the gene expression in nTPM of GPR61, GNAI3, AMIGO1, and ATXN7L2 gene expression the hypothalamic the arcuate nucleus (ARC), dorsomedial hypothalamic nucleus (DMH), lateral hypothalamic Area (LHA), ventromedial hypothalamic nucleus (VMH), and preoptic area (POA).

### Differential effect of the global knockout of *GPR61*, *AMIGO1*, and *ATXN7L2* on the body composition and activity in mice

We next queried the IMPC database to understand the impact of global knockout of mouse orthologues of high-priority candidate genes ^21^. The IMPC provides a comprehensive and standardized set of phenotypes, which ensures that each knockout mouse is evaluated using the same criteria. This standardization allows us to make direct comparisons between the effects of different gene knockouts and to understand how these genes may contribute to the traits of interest, such as body composition and activity level. Distinct physiological alterations were observed upon the global knockdown of *GPR61*, *AMIGO1*, and *ATXN7L2* in mice. *GPR61* deficiency (-/-) resulted in a significant increase in fat mass in both males and females (p < 0.01) (**Figure S3A**) and an increase in overall activity, as evidenced by the higher total and periphery distances in the open field test (p < 0.001 and p < 0.0001, respectively) (**Figure S3B-C)**. Conversely, *AMIGO1* deficiency (-/-) did not significantly affect body composition but did increase activity, with knockout mice showing an increase in total distance traveled (p < 0.01 for males and p < 0.001 for females) (**Figure S4**). In the *ATXN7L2* knockout mice, there was a significant reduction in body weight without a corresponding change in activity levels (**Figure S5**). Overall, these findings implicate a dual role in this region, both with sleep and adipose traits.

## DISCUSSION

In our study, we found a strong and significant genetic correlation between morningness chronotype and ease of getting up, as evidenced by a high genetic correlation score, with a high Human Genetic Evidence (HuGE) score for both traits. Our focus on the orphan G protein-coupled receptor GPR61 suggested it may be a high-priority candidate gene for morningness and ease of getting up in the morning. Our computational approach revealed that the lead variant, rs12044778 within *GPR61*, is a key pleiotropic variant strongly linked to both traits, as affirmed by our colocalization analysis with a posterior probability of 0.9757. Moreover, the association of rs12044778 with other circadian traits and its regulatory influence on expression patterns of nearby genes (*GPR61*, *AMIGO1,* and *ATXN7L2*), particularly in the brain and subcutaneous adipose tissues, were significant. These findings were further supported by physiological alterations observed in *GPR61*, *AMIGO1*, and *ATXN7L2* knockout mice with substantial impact on body composition and activity. Our computational approach to genetic and functional analysis at this locus using existing datasets offers a deeper understanding of the intricate relationship between morningness chronotype, ease of getting up, objective measures of sleep timing, and putative shared underlying genetic mechanisms. Our integrative approach on the lead variant rs12044778 in *GPR61* highlights the roles of three previously poorly understudied genes: *GPR61*, *ATXN7L2,* and *AMIGO1* in circadian rhythms.

GPR61 is an orphan G protein-coupled receptor (GPCR), a member of the GPCR Class A family, known for its critical roles in signal transduction processes across cell membranes. Although GPR61’s endogenous ligand remains unidentified, it shares structural similarities with other biogenic amine GPCRs involved in neurological and metabolic regulation ^27,28^. This receptor is primarily expressed in brain regions associated with behavioral and metabolic regulation ^28^, suggesting its potential role in modulating circadian and metabolic functions. Given that GPCRs are often targeted in drug discovery due to their extensive involvement in physiological processes, GPR61 presents a promising area for further investigation, particularly in the context of circadian preference and metabolic regulation. A study that used an integrative framework to prioritize genes in more than 500 loci associated with body mass index pinpointed *GPR61* as among 292 high-scoring genes from 264 loci, confirming its role in obesity in humans ^29^. Moreover, a study from Nambu et al. ^25^ adds an important layer to our understanding of GPR61’s role in metabolic processes through a GPR61-deficient mouse model generated via the insertion of the lacZ gene into the GPR61’s coding sequence. This study complements our genetic findings because the GPR61-deficient mice exhibited marked hyperphagia and heavier body weight than wild-type mice, which was accompanied by increased fat weight, liver weight, and liver triglyceride level. Moreover, this was confirmed through the IMPC mouse model, in which global removal of GPR61 using the CRISPR-cas9 system also significantly increased the fat mass in both males and females. Overall, these experimental insights into GPR61’s role in metabolism further strengthen the hypothesis that GPR61 could be a target for treating disorders that involve disruptions in both circadian rhythms and adiposity. This is particularly noteworthy given the recent cryo-EM structural characterization of GPR61 protein by Lees et al. ^27^, which will open avenues for the identification and/or design of new GPR61-specific ligands. The need for therapeutic interventions targeting GPR61 in circadian and metabolic disorders is underscored in this study by its genetic significance in circadian preferences and the observed metabolic alterations in the GPR61 mice models.

The adhesion protein AMIGO1 (Adhesion Molecule with Ig-Like Domain 1) plays a significant role in modulating the function of Kv2 voltage-gated potassium channels, which are crucial for maintaining neuronal excitability, signaling, and overall brain function ^30^. Through its interaction with these channels, AMIGO1 directly influences the transmission of electrical signals in neurons, a process that underpins complex behaviors and physiological functions ^31,32^. This modulation of Kv2 channels by AMIGO1 is essential not only for synaptic function but also for cellular communication within neural circuits ^31,33^. Given that AMIGO1 contributes to neuronal excitability and plasticity, it may have important implications for neural timing and rhythm, processes intimately connected to circadian regulation. AMIGO1’s ability to function as both a channel modulator and an adhesion molecule suggests that it may impact circadian control mechanisms, potentially affecting sleep-wake cycles and related neurological conditions. These dual roles make AMIGO1 a compelling target for further investigation in both circadian biology and neural health, as it may influence the delicate balance of excitability and stability required for daily rhythms and behaviors.

The gene *ATXN7L2* (*Ataxin 7 Like 2*), while less extensively studied, has shown associations with neurodegenerative processes, most notably through its link to spinocerebellar ataxia type 7 (SCA7), a severe and progressive neurological disorder characterized by motor coordination loss, retinal degeneration, and cognitive decline ^34^. Though its functions are not fully elucidated, ATXN7L2 is thought to be involved in maintaining neural homeostasis, potentially by supporting cellular structures that prevent neurodegeneration. The gene’s connection to SCA7 highlights a possible role in stabilizing neural circuitry, particularly within cerebellar and motor pathways ^34^. Additionally, ATXN7L2 may play a role in circadian biology by supporting the stability of neurons responsible for timing and rhythm. The link to neurodegeneration raises questions about the role of ATXN7L2 in neuroprotective mechanisms, suggesting it could influence not only circadian function but also the resilience of neural networks. As such, understanding ATXN7L2’s role in both disease and circadian regulation could provide valuable insights into potential therapeutic targets for neurodegenerative diseases and circadian-related disorders.

Our study provides further insight into the lead colocalized variant rs12044778 at this locus. Its association with additional circadian traits and its tissue-specific regulatory role in gene expression, particularly in the brain and subcutaneous adipose tissues, highlights the variant’s broader influence on circadian and metabolic functions. Our study’s use of Hi-C chromatin interaction loop analysis from adipose-progenitor mesenchymal stem cells provided novel insights into the regulatory mechanisms of rs12044778’s GPR*61*, revealing that rs12044778 interacts with a network of nearby genes, suggesting a complex regulatory landscape where rs12044778 potentially influences multiple genetic pathways. Furthermore, we demonstrated that rs12044778 is predicted to alter the binding of transcription factors EGR1 and NR2F2, providing further evidence that rs12044778 may exert its influence on morningness genotype and ease of getting up by modulating the expression of *GPR61* and its nearby genes.

Moreover, our tissue-specific eQTL analysis provided additional evidence of rs12044778’s functional impact. In nerve tibial tissue, rs12044778 allele A significantly decreased *GPR61* gene expression while it increased *AMIGO1* expression in subcutaneous adipose tissue and *ATXN7L2* expression in the cerebellar hemispheres. The colocalization of genetic association signals and eQTL signals further refined our understanding of the genetic landscape influenced by rs12044778. Notably, rs1434285, in linkage disequilibrium with rs12044778, colocalized with AMIGO1 expression in subcutaneous adipose tissue and GNAI3 expression in the cerebellum, illustrating the specific genetic interactions that contribute to morningness chronotype and ease of getting up on the morning. This differential expression pattern in tissues underlines the variant’s systemic influence. The exact nature of this relationship is complex and may involve multiple pathways, including sensory input, modulation of environmental cues, and systemic genetic influences.

Although we present compelling evidence for the involvement of rs12044778 in regulating circadian and metabolic functions, our study’s limitations include the need for replicability and generalizability of its findings across broader and more diverse populations to confirm their applicability across different genetic backgrounds and environmental contexts. Additionally, while our computational analyses suggest potential mechanisms for rs12044778’s effects on gene expression, direct functional assays in human tissues are required to fully understand the variant’s molecular impact. Furthermore, we recognize that this variant alone may not explain the full impact of the locus on the circadian preference phenotypes.

Overall, our comprehensive integrative genetic and functional analysis offers a deeper understanding of the GPR61 locus and the complex relationship between morningness chronotype, ease of getting up, and their underlying genetic mechanisms.

## ACKNOWLEDGMENTS

This research has been conducted using the UK Biobank Resource (UK Biobank application number 6818). We would like to thank the UKB study participants and researchers.

C.T. is supported by the BWF G-1022367 and NIH 5T32HL007609-35 and R01HL146751 (RS). J.W. is supported by the NIH R01-HL-127146. H.T. is supported by the NIH UL1TR002378 and NIH P50HL117929. H.M.O. and R.S. are supported by R01AI170850, and R.S. is further supported by R01HL146751.

## Author contributions

This study was designed by C.T. and R.S., and they participated in the acquisition, analysis, and/or interpretation of data. C.T. wrote the manuscript, and all co-authors reviewed and edited the manuscript before approving its submission.

