## Supplementary material for "An integrative approach prioritizes the orphan GPR61 genomic region in tissue-specific regulation of chronotype": Figure S

**Supplemental Figures**

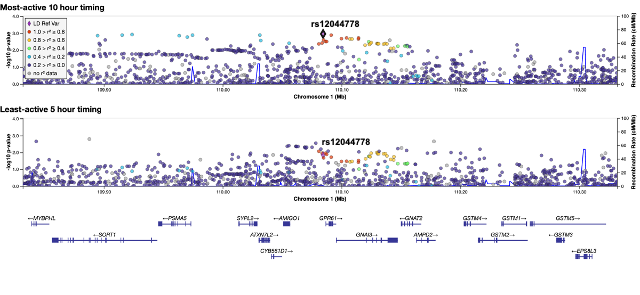

**Figure S1: Regional Plot of GPR61 loci in significant circadian traits of waking up GWAS.** Locuszoom was used to generate the regional plot of colocalized *GPR61* loci with GWAS significances (P-value = 0.005 {0.05/10 Circadian Phenotypes}) for circadian traits: Most-active 10-hour timing and least-active 5-hour timing. The Lead variant rs12044778 (purple diamond) is in LD (EUR Ancestry) with other variants. The significance threshold of 0.005 (0.05/10 Circadian Phenotypes)

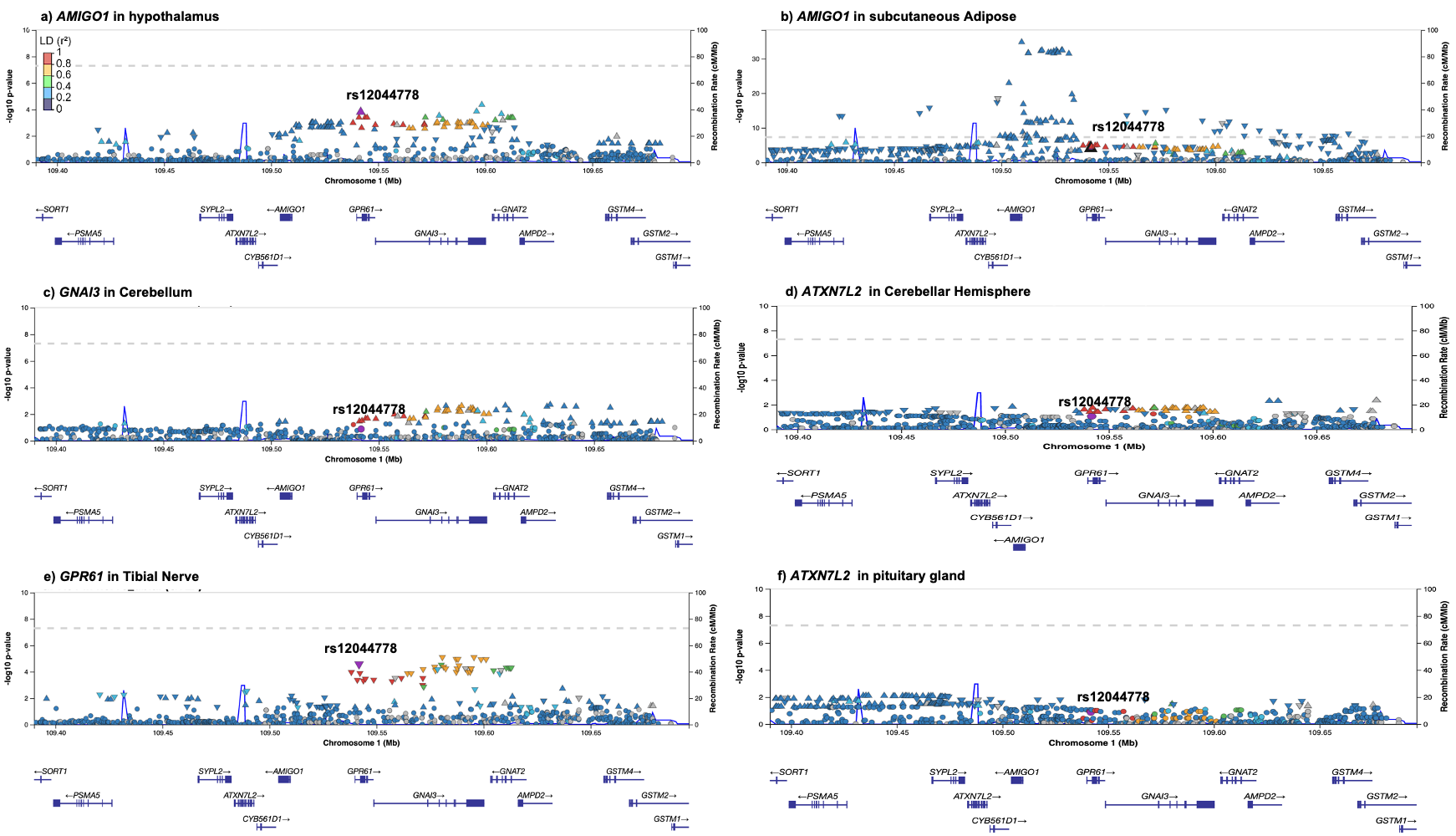

**Figure S2: Colocalization of tissue-specific eQTL of lead variant rs12044778 from morning chronotype and ease of waking up GWAS analysis**. Colocalization of single-tissue cis-eQTLs with chronotype and ease of waking up. Plots of the chromosome 1 locus rs12044778 depict the colocalized allele significantly increases *AMIGO1* expression in (**a**) the hypothalamus (n=202); (**b**) subcutaneous adipose (*n* = 663); it increases (**c**) *GNAI3* expression the cerebellum (n=241); *ATXN7L2* expression in cerebellar hemisphere (n=175); (**e**) the colocalized allele rs12044778 significantly decrease *GPR61* signal in the tibial nerve (n=619); *ATXN7L2* expression in pituitary gland (n=283) The P-value thresholds are in **Table S3**. The lead variant rs12044778 is purple in color and is the lead variant (*r^2^* > 0.8). Plots genomic region are chr1:109389996-109695867.

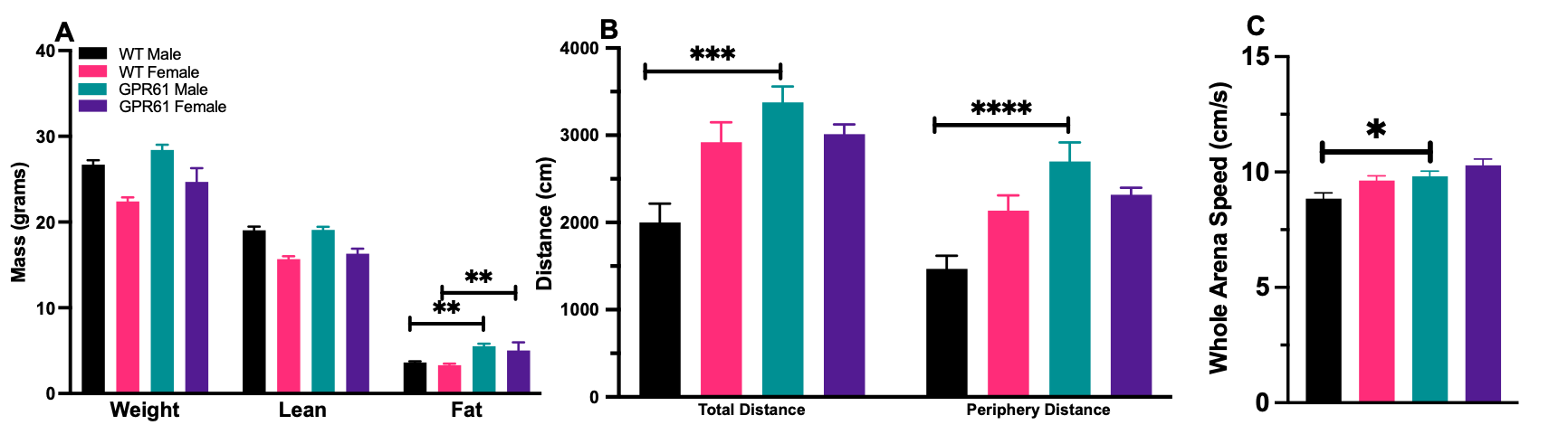

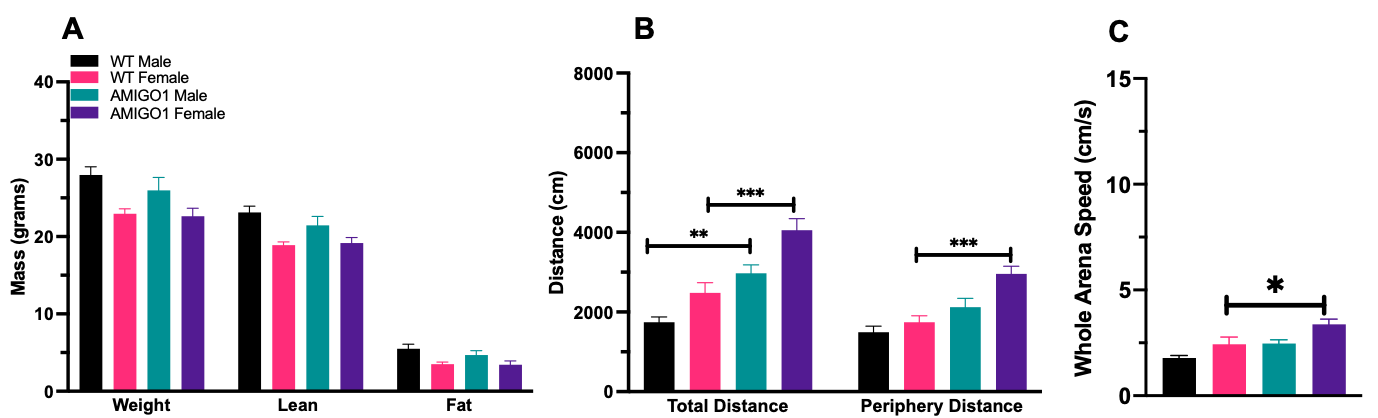
**Figure S3: The removal of *GPR61* increases the mice's fat mass and activity.** The mice's body composition **(A)** was measured in grams. The open field test for activity was measured in **(B)** distance (cm) and **(C)** whole arena speed (cm/s) The results are expressed in mean ± SEM (n=7-10/genotype-sex; two-way ANOVA followed by Tukey post hoc test, *P < .05, **P < .01, ***P < .001, and ****P < .0001).

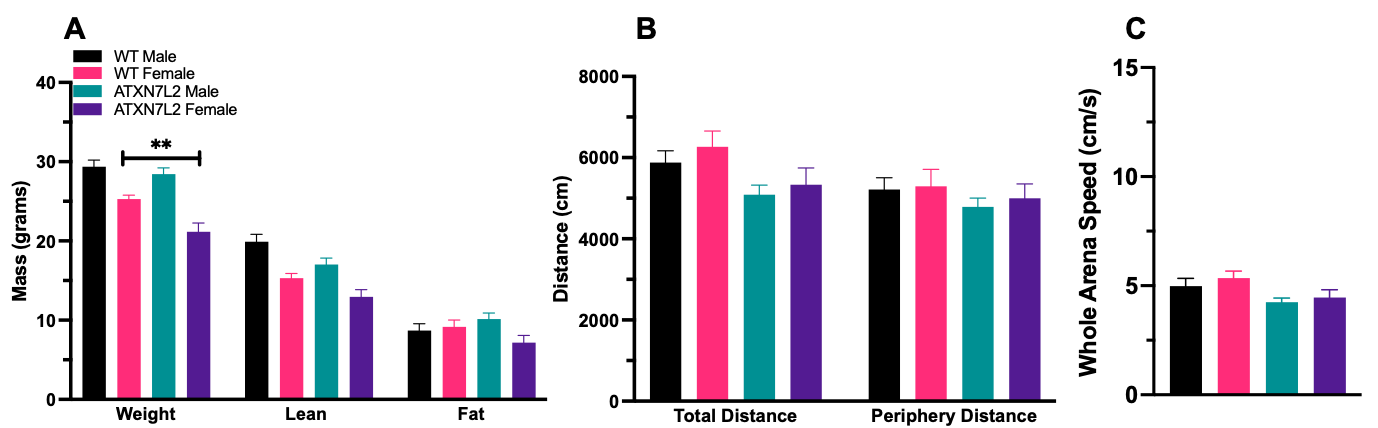
**Figure S4: The removal of *AMIGO1* increases the mice's activity.** The mice's body composition **(A)** was measured in grams. The open field test for activity was measured in **(B)** distance (cm) and **(C)** whole arena speed (cm/s) The results are expressed in mean ± SEM (n=7-10/genotype-sex; two-way ANOVA followed by Tukey post hoc test, *P < .05, **P < .01, ***P < .001, and ****P < .0001).

**Figure S5: The removal of *ATXN7L2* decreases the mice's body weight.** The mice's body composition **(A)** was measured in grams. The open field test for activity was measured in **(B)** distance (cm) and **(C)** whole arena speed (cm/s) The results are expressed in mean ± SEM (n=7-10/genotype-sex; two-way ANOVA followed by Tukey post hoc test, *P < .05, **P < .01, ***P < .001, and ****P < .0001).
